# SHEST: Single-cell-level artificial intelligence from haematoxylin and eosin morphology for cell type prediction and spatial transcriptomics reconstruction

**DOI:** 10.1101/2025.11.19.689364

**Authors:** Hoyeon Jeong, Junghan Oh, Donggeon Lee, Jae Hwan Kang, Yoon-La Choi

## Abstract

A comprehensive understanding of cancer progression requires integrating tissue morphology with spatial molecular profiles. We present SHEST, a multi-task profiling framework that leverages haematoxylin and eosin morphology to predict cellular composition and reconstruct spatial gene expression at single-cell resolution. SHEST employs a quadruple-tile input capturing nuclear and contextual information, combined with a neighbourhood-informed clustering algorithm to filter ambiguous cellular signals. It comprises a shared morphological encoder with two task-specific heads: a classifier for cell type prediction and a reconstructor for gene expression. Multi-task optimisation uses cross-entropy and zero-inflated negative binomial losses, specifically addressing the sparsity of spatial transcriptomic data. Evaluation on human lung adenocarcinoma datasets demonstrated high accuracy for the principal reciprocal constituents of the tumour–immune axis (*F*_1_: 0.97 for tumour cells and 0.91 for lymphocytes). External validation confirmed its generalisability, revealing alveolar cells and their early neoplastic transitions. Reconstructed gene expression reproduced spatially resolved, cell-type-specific marker patterns—*EPCAM* in tumour cells, *LTBP2* in fibroblasts, and *CD3E* in lymphocytes—recovering biologically coherent transcriptional architecture. SHEST also preserved distance-dependent spatial relationships and gene-level autocorrelation, reflecting the multicellular niche structure of the tumour microenvironment. By unifying cell type identification, gene expression reconstruction, and spatial mapping within a single interpretable framework, SHEST provides a synergistic and cost-efficient bridge between histopathology and spatial transcriptomics. This approach facilitates comprehensive tissue characterisation and forms a foundation for precision oncology through spatially informed, cell-level insights into tumour–immune ecosystems.

## 1 Introduction

Achieving a comprehensive understanding of cancer progression demands the simultaneous examination of tissue morphological organisation and the precise spatial distribution of functional molecular signatures [1, 2]. Over the past century, cancer diagnosis and interpretation have primarily depended on morphological examination of tissue sections, most commonly through haematoxylin and eosin (H&E) staining [3]. This classical histopathological approach, which visualises the structure of cell nuclei and cytoplasm, provides essential information about tissue architecture and disease progression. In recent decades, advances in molecular biology and next-generation sequencing technologies have transformed the understanding of cellular function by revealing that the state of a cell is defined by its gene expression profile [4]. The quantitative study of complex tissue systems, such as the tumour microenvironment (TME), now depends on the capacity to link cellular morphology with molecular function [5, 6]. Establishing this correlation is fundamental for developing precision medicine and gaining deeper insights into disease mechanisms.

H&E staining produces morphological contrast by colouring nuclei blue with haematoxylin and cytoplasm or extracellular components pink with eosin. The development of digital pathology (DP), which converts glass slides into high-resolution whole-slide images (WSIs), has enabled faster and more consistent diagnostic processes. This digital transformation has also facilitated quantitative analysis, improving reproducibility compared to traditional, observer-dependent assessments [7]. In parallel, molecular profiling technologies have progressed rapidly. The emergence of single-cell RNA sequencing (scRNA-seq) has been followed by the introduction of spatial transcriptomics (ST) technologies [8]. These methods measure RNA expression while preserving spatial context, allowing researchers to map gene expression patterns, infer cell type distributions, and analyse cell-cell interactions within the tissue architecture. Such analyses are particularly valuable for studying cancer biology and immune regulation within the TME [9].

DP and ST exhibit complementary strengths and limitations. DP provides high-resolution morphological context that is indispensable for histological interpretation and cancer grading. However, it lacks molecular information and is influenced by inter-observer variability. In contrast, ST yields spatially resolved molecular profiles that capture the functional state of cells, yet it is limited by relatively coarse visual resolution and high experimental costs. Integrating these two modalities enables a comprehensive approach that combines morphological context with molecular characterisation at the single-cell level [10].

### 1.1 Related work

Research linking H&E histology with molecular gene expression data can be broadly categorised into two main directions: morphological classification of individual cells or nuclei, and computational inference of spatial molecular profiles from histological images.

Accurate single-cell or nuclei segmentation and classification are essential prerequisites for spatial analysis at cellular resolution. One of the representative methods in this area is HoVer-Net [11], which introduced a dual-task network capable of performing nuclei instance segmentation and classification simultaneously by predicting horizontal and vertical distance maps to separate clustered nuclei. HD-Yolo [12] adopted a YOLO [13]-based framework for nuclei localisation and segmentation, reporting that morphological features extracted from the model were correlated with patient survival in breast cancer. CellViT [14] applied a Vision Transformer (ViT) [15] architecture to enable parallel processing and local feature extraction through attention mechanisms. While these methods advanced morphological analysis, the overall classification accuracy remains still low, and most models have not undergone comprehensive external validation.

A complementary research stream focuses on predicting molecular information directly from H&E images. Most prior studies, such as mclSTExp [16] and TCGN [17], have been developed for sequencing-based ST platforms such as Visium, where each spot corresponds to an area of approximately 55 µm that contains multiple cells rather than a single cell. These models employ a regression head trained with the mean squared error (MSE) loss function to infer gene expression levels at the Visium spot level. In addition, two studies, Hist2Gene [18] and THItoGene [19], which modelled local features using convolution, global features using ViT, and connectivity features using GNN on breast cancer and squamous cell carcinoma tissues, are also noteworthy. Despite their methodological diversity, all the aforementioned models share a fundamental limitation: they are mostly trained using publicly available Visium data from breast cancer tissue, which represents unpurified or unseparated profiles where multiple cells are mixed within a single spot. That is, the prediction outcome represents an average or aggregate of the multiple cells within the spot, which inherently precludes true single-cell-level analysis.

The recent development of imaging-based ST technologies, such as Xenium, enables molecular profiling at single-cell resolution. Xenium supports 5K gene panels and provide aligned H&E images. Utilising this advancement, SPiRiT [20] introduced a framework employing a ViT backbone to map single-cell H&E image tiles to corresponding cellular gene expression profiles. This model demonstrated the feasibility of inferring cell-level molecular signatures from cellular morphology. However, SPiRiT still uses the MSE loss function for gene expression regression, which is not optimal for modelling zero-inflation characteristic of RNA sequencing data. Furthermore, the limited external validation restricts confidence in the model’s generalisability across diverse tissue types and cohorts.

Despite these pioneering efforts, a methodological discontinuity remains in simultaneously and coherently addressing the multifaceted nature of cellular state, encompassing both morphology and molecular function. The development of a unified framework that jointly performs cell-level morphological encoding, cell type classification, and gene expression reconstruction directly from H&E images is therefore presented as the necessary next step to advance single-cell spatial inference.

### 1.2 Proposed framework: SHEST

SHEST (Single-cell-level artificial intelligence from H&E morphology for ST reconstruction and cell type prediction) is a multi-task profiling framework that integrates morphological and molecular information at single-cell resolution (Figure 1). The framework aligns morphological features extracted from H&E images with molecular features obtained from ST on shared spatial coordinates, enabling a direct correspondence between each cell’s morphology and its gene expression profile (Figure 1A). Through this, SHEST simultaneously performs two learning tasks. The first is the classification of cell types based on morphological context. The second is the reconstruction of single-cell gene expression from image-derived embeddings (Figure 1B and C). This joint optimisation enables the model to share representations across tasks, ensuring consistency between morphological classification and molecular prediction. Overall, SHEST offers an integrated computational approach for single-cell molecular profiling from H&E images, establishing a direct and interpretable connection between tissue morphology and gene expression (Figure 1E).

**Figure 1:**
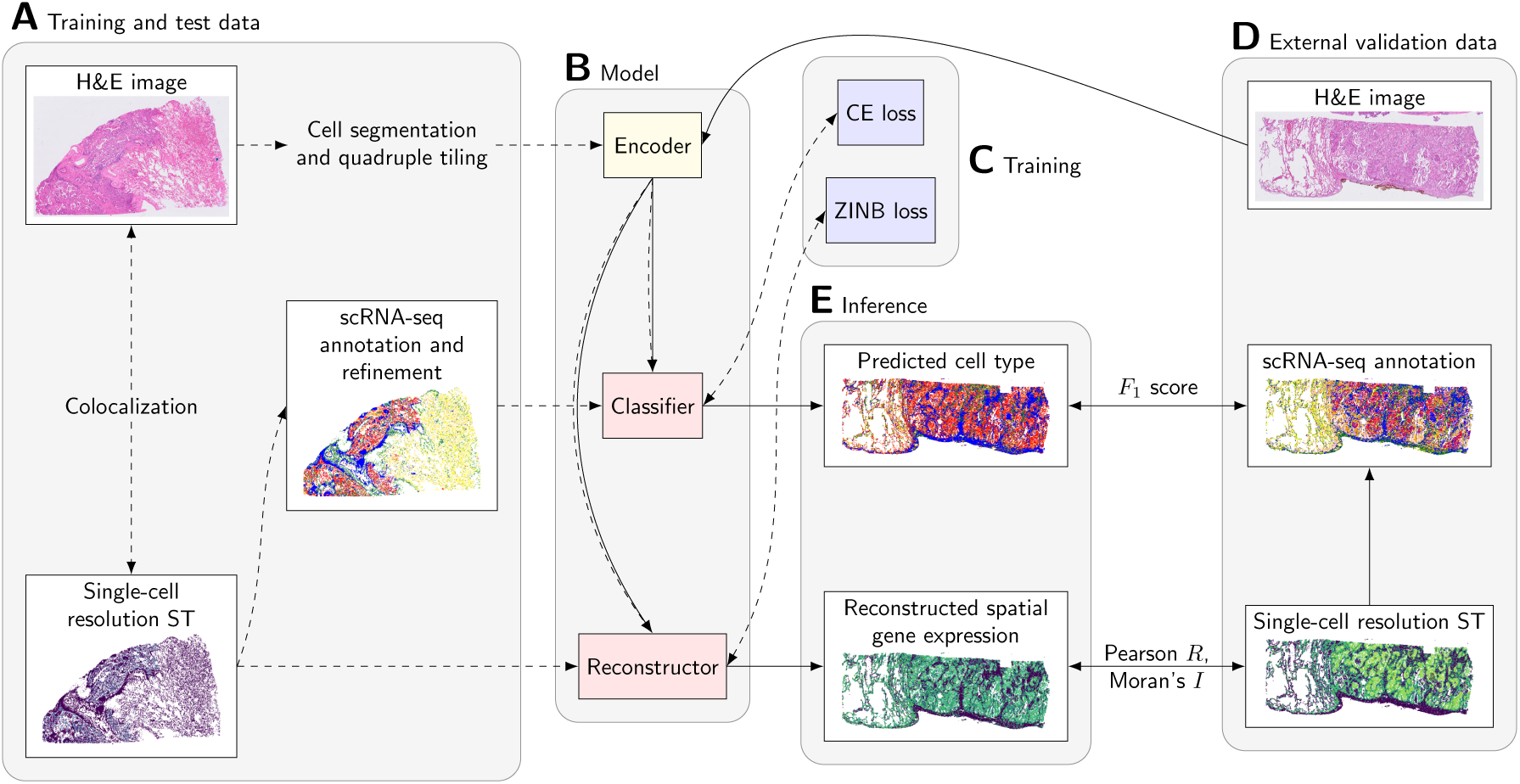
Overview of the SHEST framework and its workflow. **A** Training and test data: H&E images are processed through cell segmentation and quadruple tiling to generate per-cell image tiles. Single-cell–resolution spatial transcriptomics (ST) data are annotated with cell types using reference single-cell RNA sequencing (scRNA-seq) profiles. These multimodal datasets are spatially aligned to construct paired image–transcriptome–label sets for supervised training. **B** Model: A shared encoder extracts morphological embeddings from the input image tiles. Two task-specific heads operate on these embeddings: a classifier that predicts cell types and a reconstructor that generates reconstructed spatial gene expression profiles for each cell. **C** Training: Multi-task learning is supervised by two loss functions — cross-entropy (CE) loss for classification and zero-inflated negative binomial (ZINB) loss for gene expression reconstruction. Dashed arrows in the diagram indicate training-time data flow and supervision paths connecting the ground-truth labels and measured expression to their respective losses. **D** External validation data: Independent H&E images, single-cell–resolution ST, and scRNA-seq annotation datasets are used to assess generalisation performance. **E** Inference: The trained model generates cell-type predictions and reconstructs spatial gene expression maps for each localised cell. Solid arrows indicate the forward data flow through the encoder and both task heads. Model performance is quantitatively evaluated using F_1_ scores for classification accuracy and Pearson’s R and Moran’s I for assessing expression-level agreement and spatial autocorrelation.

## 2 Materials and methods

The SHEST framework was implemented in Python 3.10 using PyTorch 2.6.0+cu124 [21] for modelling, and SpatialData 0.3.0 [22] and Scanpy 1.10.4 [23] for processing Xenium data.

### 2.1 Data acquisition

The study utilises ST data coupled with matched H&E WSIs from lung adenocarcinoma (LUAD) samples, a highly representative and clinically significant tumour type. Detailed information regarding the datasets is summarised in Table 1.

**Table 1:**
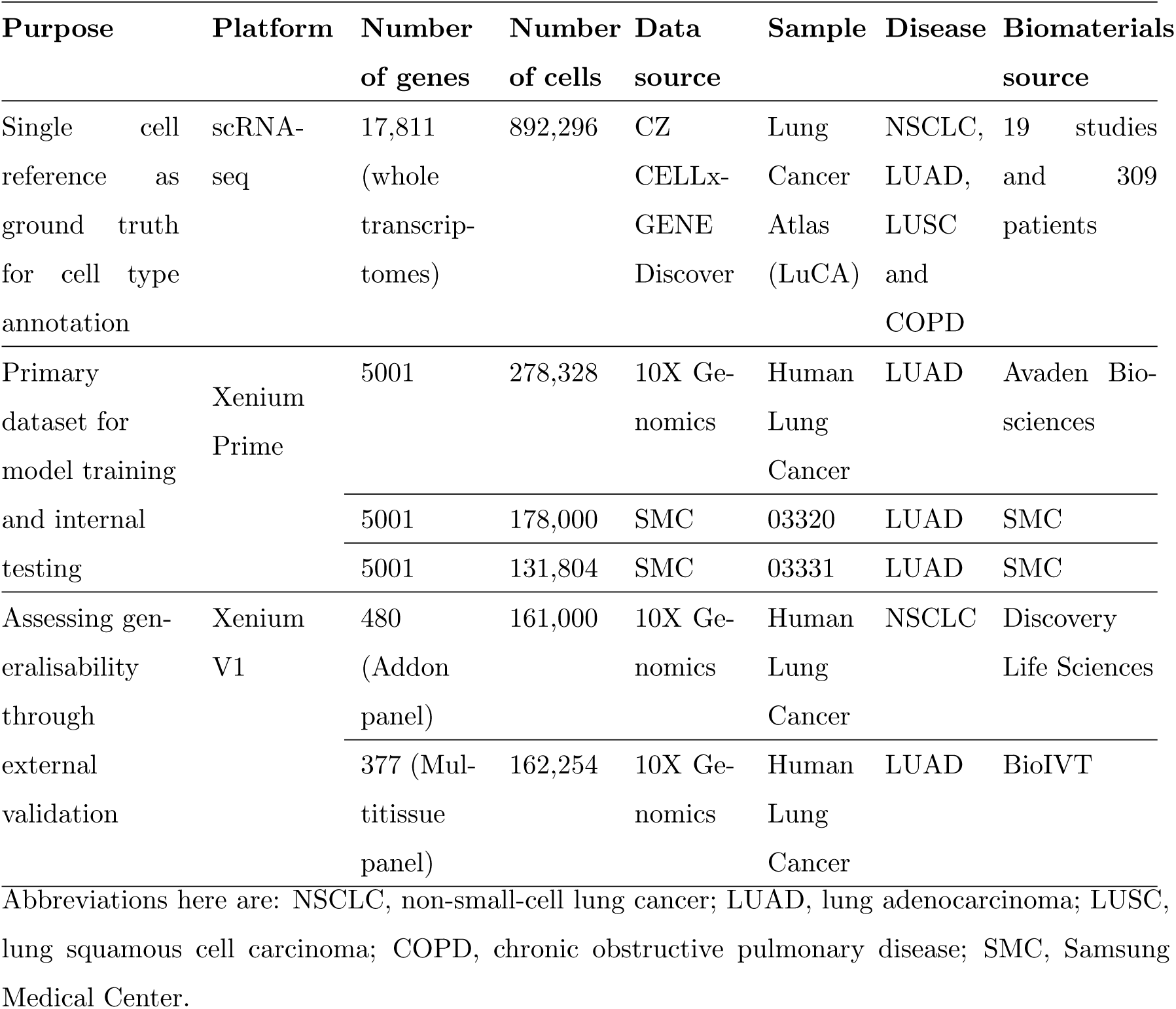
Overview of datasets used for SHEST model development and validation.

The foundation for our ground truth cell type labels is the comprehensive CZ CELLxGENE Discover - Human Lung Cancer Atlas (LuCA) [24]. The primary dataset used for training and internal testing of the SHEST model is the 10X Genomics Xenium Prime Human Lung Cancer dataset (Figure 1A). For external validation and assessment of the model’s generalisability, we employed two of the earlier 10X Genomics Xenium V1 Human Lung Cancer datasets, which are characterised by a lower gene count panel compared to the Xenium Prime panel (Figure 1D). All H&E slides were scanned at a magnification of ×40 objective.

### 2.2 Cell type annotation

Cell identities were initially annotated on the Xenium Prime data using the TACCO [25] package against the LuCA single-cell references (Figure 1A). Following this initial computational step, the resulting annotations were subjected to thorough pathologist (Y.C.) review for refinement and validation. This review utilised the rich morphological information provided by the H&E image, integrating not only the distinct cellular morphology but also the spatial and locational context. Specifically, the pathologist assessed critical biological patterns, such as tumour cells lining the boundary, the floating status of macrophages, the spatial relationship between endothelial cells and neighbouring erythrocytes (RBCs), and the formation of tertiary lymphoid structures (TLS) by lymphocytes. These validated and detailed subtypes were then systematically grouped into higher-level cell types, as summarised in Supplementary Table S1, to facilitate robust classification and downstream analysis within the SHEST framework.

### 2.3 Image preparation for encoding

The preparation of H&E image data for morphological encoding involves sequential steps of spatial alignment, cell filtering, patch extraction, and colour standardisation. First, precise spatial co-registration between the H&E image coordinates and the ST coordinate system is established (Figure 1A). This alignment is achieved by scaling the raw image coordinates according to the known pixel size and subsequently applying an inverse affine transformation. Following alignment, individual cells, identified via segmentation, are filtered based on their bounding box size. Only cells with dimensions within the range of 2 µm × 2 µm to 16 µm × 16 µm are retained (Figure 2A); excessively small or large cells are excluded to prevent misclassification stemming from insufficient or distorted morphological information. Next, we constructed the local morphological context for each retained cell using four concatenated tiles: the whole cell, the nucleus-masked cell, the haematoxylin extraction, and the nucleus-masked haematoxylin extraction (Figure 2B-G). This process generates a set of image patches designed to encompass both the local cellular environment and focused nuclear features. This strategy ensures the encoder receives comprehensive morphological data, prioritising nuclear-specific details while reducing extraneous peripheral context. Finally, the extracted H&E image windows are standardised to fulfil the input requirements of the H-optimus-0 [26] encoder. All patches are resized to a fixed resolution of 224 × 224 pixels, after which the pixel values are normalised by dividing the image tensor by 255. A colour normalisation process is then applied for RGB standardisation, using the pre-calculated mean (0.707223, 0.578729, 0.703617) and standard deviation (0.211883, 0.230117, 0.177517) specific to the H-optimus-0 model.

**Figure 2:**
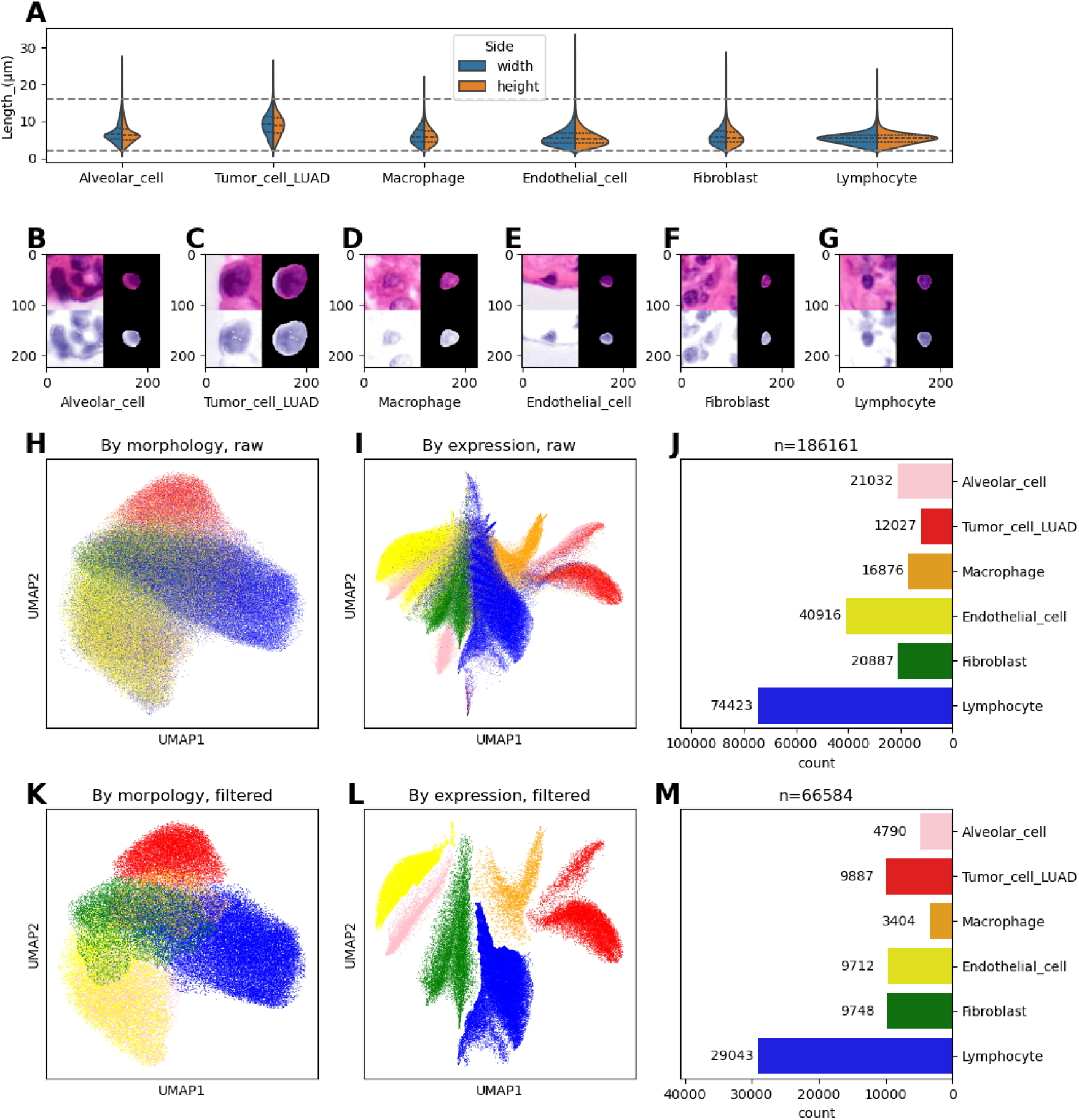
Morphological and molecular characteristics of cell types before and after annotation refinement for the 10X Xenium Prime Human Lung Cancer sample. Panel **A** shows the distribution of cell size parameters (width and height in µm) as violin plots for the six cell types. Panels **B-G** show morphological examples of six cell types, with each panel displaying a 224 × 224 pixel image patch corresponding to a single cell: Alveolar_cell (**B**), Tumor_cell_LUAD (**C**), Macrophage (**D**), Endothelial_cell (**E**), Fibroblast (**F**), and Lymphocyte (**G**). Quadruple tiles with nucleus extraction and haematoxylin extraction are used for each patch. Panels **H**-**J** present the raw data 25 before annotation refinement: **H** UMAP visualisation based on raw gene expression profiles, **I** UMAP visualisation based on H-optimus-0 morphological embeddings, and **J** cell count distribution across the six cell types. Panels **K**-**L** show the data after annotation refinement: **K** UMAP visualisation of the curated population based on gene expression, **L** UMAP visualisation based on morphological features, and **M** cell count distribution of the high-purity, curated dataset.

### 2.4 Annotation refinement for high-fidelity training

To ensure the SHEST model is trained on data with high confidence and distinct molecular-morphological correspondence, we implemented an annotation refinement step prior to model training (Algorithm 1). This procedure maximises the clarity of learnt features by filtering out cells situated in low-density regions or near cluster boundaries within the latent space, where cell signatures are often ambiguous or mixed. The refinement process operates on low-dimensional representations derived from both modalities. First, the gene expression profile and the corresponding morphological features of all cells are projected into separate low-dimensional UMAP [27] spaces, creating distinct embeddings that represent molecular and morphological cell clustering, respectively. Starting with the cell type class that has the smallest cell count (to prevent disproportionate pruning of minority populations), we iteratively curate the dataset using the DBSCAN [28] clustering algorithm. Only the single dominant DBSCAN cluster for the target cell type is retained. Concurrently, any cells of other types that neighbour this dominant cluster within the defined *k*-nearest neighbours are effectively marked for removal, preventing the inclusion of cells at ambiguous class boundaries. This systematic curation, performed on both the expression and morphological feature spaces, efficiently removes cells that exhibit ambiguous or contradictory signatures, ultimately retaining only those cells that are common to both modalities.

Morphological analysis of the six cell types reveals significant size variations that serve as distinguishing features (Figure 2G). The Tumor_cell_LUAD population exhibits the largest relative size among all cell types, characterised by nuclear pleomorphism [29]. Conversely, the Lymphocytes are characterised by consistently displaying the smallest physical dimensions and large cell numbers. Beyond size, visualisation of the data in the low-dimensional feature space before refinement (Figure 2H and 2I) shows considerable overlap, indicating ambiguity in raw cell classification based on either expression or morphology. Applying the annotation refinement algorithm resulted in a reduction in the total cell count by effectively removing low-confidence and boundary cells (Figure 2K and 2L). The UMAP visualisations of the refined dataset demonstrate markedly enhanced cluster purity and more distinct cluster boundaries in both the expression and morphological feature spaces.

**Algorithm 1** Representative cell selection with neighbour-aware clustering

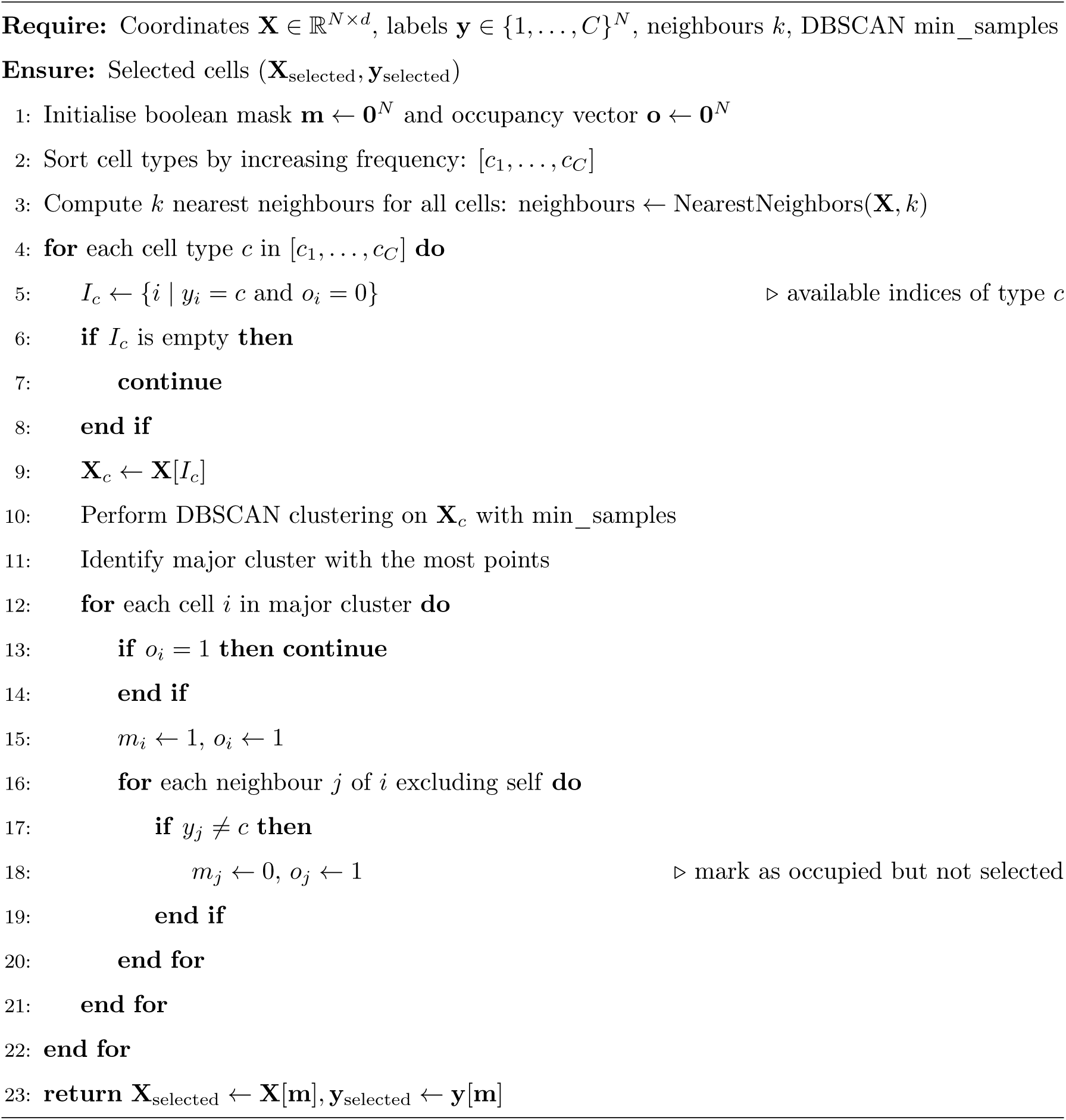

### 2.5 Data augmentation and split

To mitigate the effects of class imbalance inherent in biological samples, a data augmentation strategy is implemented during the dataset-loading phase. This procedure effectively mimics random oversampling by applying geometric transformations to the H&E image patches. Specifically, the augmentation includes random rotations (one of 0*^◦^,* 90*^◦^,* 180*^◦^,* 270*^◦^*) and horizontal or vertical flipping. By generating multiple transformed copies of minority class samples, this strategy balances the number of samples across all cell classes, thereby promoting robust and unbiased learning by the subsequent model layers.

The final dataset was divided into training and testing splits (80:20) using a fixed generator.

### 2.6 Architecture of the SHEST Model

The SHEST model is implemented as a unified multi-task learning framework that jointly reconstructs single-cell gene expression and classifies cell types from H&E-stained image patches. The architecture is composed of three integrated modules—an encoder, a classifier, and a reconstructor—that share a common latent embedding to couple morphological and molecular representations (Figure 1B and 3). This multi-head design thereby moves beyond a mere juxtaposition of tasks, enabling complementary learning dynamics that enhance interpretability by aligning morphological cues with their underlying molecular correlates.

**Figure 3:**
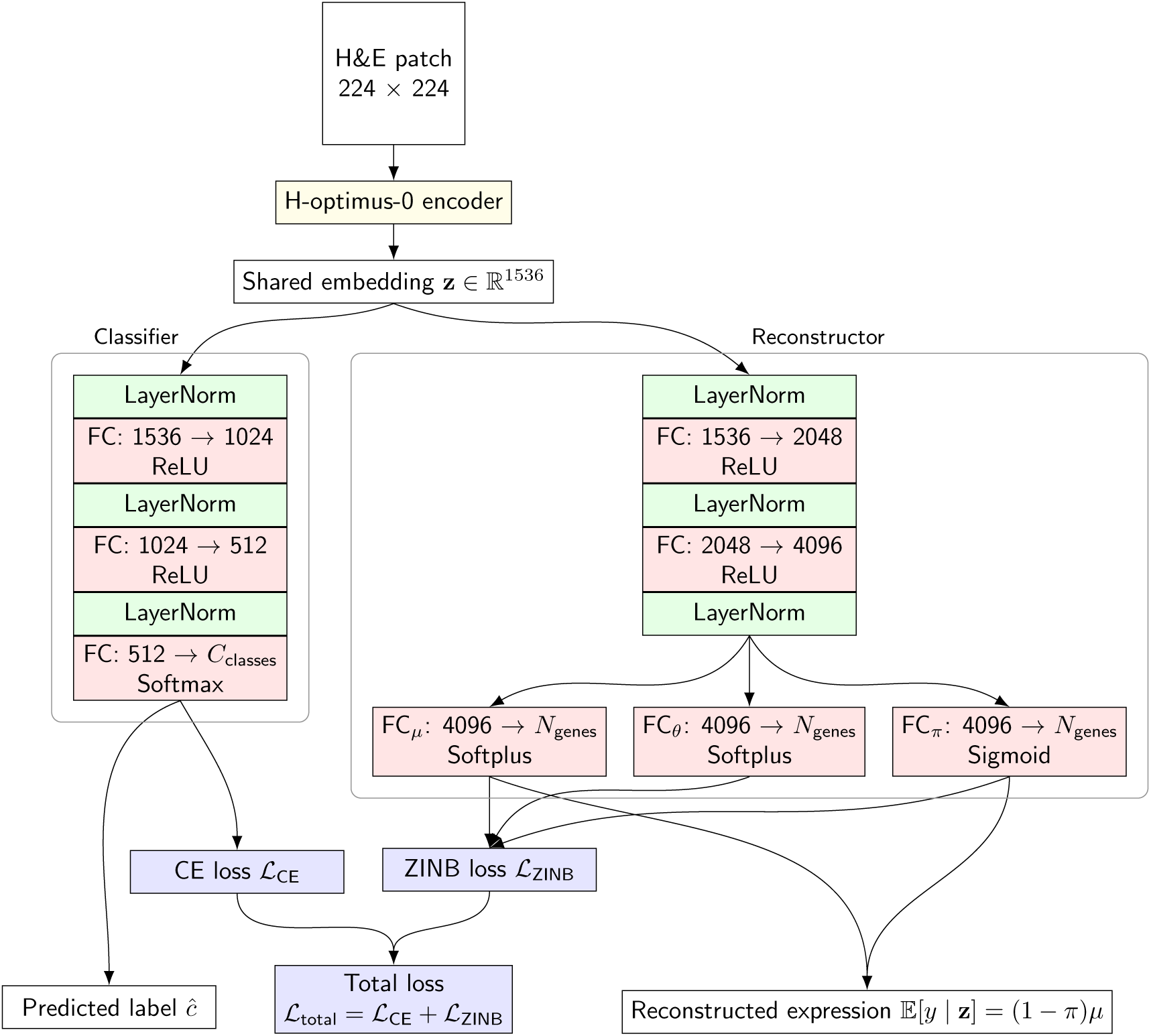
Overview of the SHEST model architecture. The model jointly classifies cell types and reconstructs single-cell gene expression from H&E-stained image patches. An input patch is encoded by a pretrained H-optimus-0 encoder into a shared embedding **z**. The Classifier branch predicts the refined cell identity through multiple fully connected (FC) layers with Layer Normalisation optimised by the cross-entropy loss L_CE_. The Reconstructor branch predicts the parameters µ, θ, and π of the zero-inflated negative binomial (ZINB) distribution through separate FC heads and reconstructs the expected gene expression E[y | **z**], optimised by the ZINB loss L_ZINB_.

The encoder is based on the H-optimus-0 ViT backbone, which is pre-trained on a large-scale biological image dataset. During SHEST training, all encoder parameters are frozen to preserve the morphological representations learnt during pre-training. The encoder processes an input H&E image of size 224 × 224 pixels and outputs a 1536-dimensional embedding vector **z** ∈ P^1536^ that serves as the common latent representation for all downstream tasks.

The classifier receives the latent embedding **z** and predicts the refined cell identity. It is structured as a multi-layer perceptron (MLP) with Layer Normalisation (LN) [30], ReLU activation, and Dropout (0.3). The final layer outputs logits **o** for *C* high-level cell type categories:

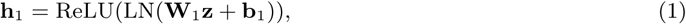

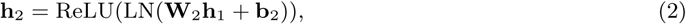

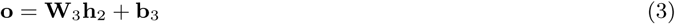

where **W***_l_* and **b***_l_* are the learnable parameters for the *l*-th layer (where *l* ∈ {1, 2, 3}), and *o_i_*is the logit corresponding to class *c_i_*. The class probability for category *c_i_* is obtained from the logits using the Softmax function:

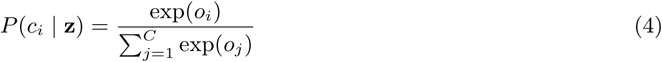

The reconstructor maps the latent embedding **z** to the predicted single-cell gene expression levels. It employs an MLP structure, generating an intermediate representation **h**_2_:

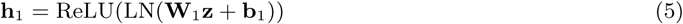

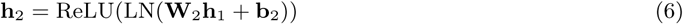

This representation **h**_2_ is then projected into three separate linear heads to estimate the parameters of the zero-inflated negative binomial (ZINB) distribution [31]—mean (*µ*), dispersion (*θ*), and zero-inflation probability (*π*):

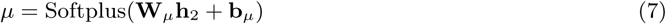

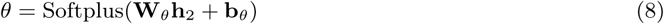

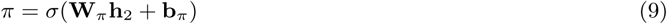

The ZINB distribution is used to model the count nature and high sparsity of single-cell expression data. The probability mass function for an observed gene count *y* = *k* given the latent embedding **z** is *P* (*y* = *k* | **z**) and is defined by its predicted parameters:

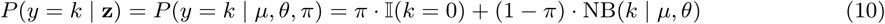

where Ι(*k* = 0) is the indicator function, and NB(*k* | *µ, θ*) is the negative binomial distribution. The reconstructed expression *y*^ is derived from the expected value E[*y* | **z**] = (1 − *π*)*µ*.

### 2.7 Loss functions and optimisation

The SHEST model is trained by jointly optimising the classification and reconstruction objectives via a multi-task loss function (Figure 1C). The classification task minimises the cross-entropy (CE) loss L_CE_, which is equivalent to the negative log-likelihood (NLL) of the true class distribution for N cells:

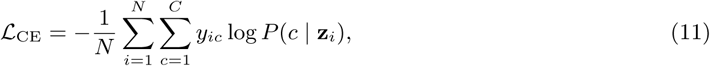

where *y_ic_* is the ground-truth indicator for class *c* of cell *i*, and *P* (*c* | **z***_i_*) is the class probability predicted by the classifier based on the embedding **z***_i_*.

The reconstruction task minimises the NLL of the ZINB distribution L_ZINB_ for the observed gene count vector **y***_i_*:

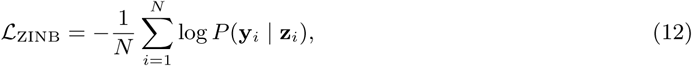

where *P* (**y***_i_* | **z***_i_*) is the likelihood of observing the ground-truth gene counts **y***_i_* under the ZINB parameters predicted from **z***_i_*. The final multi-task loss L is the unweighted sum of the two objectives:

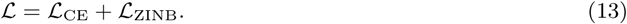

This joint optimisation ensures consistency between the predicted cell type and reconstructed gene expression. The model is optimised using the AdamW [32] optimiser with a learning rate of 0.01, trained for 40 epochs with a batch size of 1024, and utilises mixed precision training and early stopping.

### 2.8 Deterministic learning for reproducibility

To ensure the reproducibility and reliability of the SHEST framework, strict determinism was enforced across both configuration and model initialisation stages. A fixed global seed of 42 was set for NumPy and PyTorch to control stochastic operations. During data splitting, PyTorch used a fixed generator initialised with this seed. Computational determinism was further guaranteed by configuring the cuDNN [33] back-end to use deterministic algorithms and by disabling benchmarking, ensuring that identical inputs yield consistent outputs regardless of parallelisation. Consistent parameter initialisation is critical for reproducible training trajectories. All FC layers in the multi-task prediction heads, as well as any custom convolutional layers, were initialised using the Kaiming He method [34] to maintain stable gradient variance. This standardised initialisation ensures that the model starts from the same parameter state in every training run, providing reliable and repeatable results.

## 3 Results

### 3.1 SHEST enables simultaneous cell type prediction and transcriptomic profiling from H&E patches with high accuracy

The proposed dual-task model was evaluated on the test dataset derived from the 10X Xenium Prime Human Lung Cancer case to assess its capability to classify cell types and reconstruct gene expression profiles from histological image patches.

SHEST achieved high classification performance across all six major lung cancer–associated cell types (Figures 4A and 4B). The confusion matrices demonstrate that the model distinguished morphologically ambiguous but functionally distinct populations. SHEST achieved particularly high accuracy for tumour cells (*F*_1_ = 0.97) and lymphocytes (*F*_1_ = 0.91), the two pivotal cellular populations that define the tu-mour–immune landscape. In this context, the weighted *F*_1_ score is 0.87 (Figure 4C).

**Figure 4:**
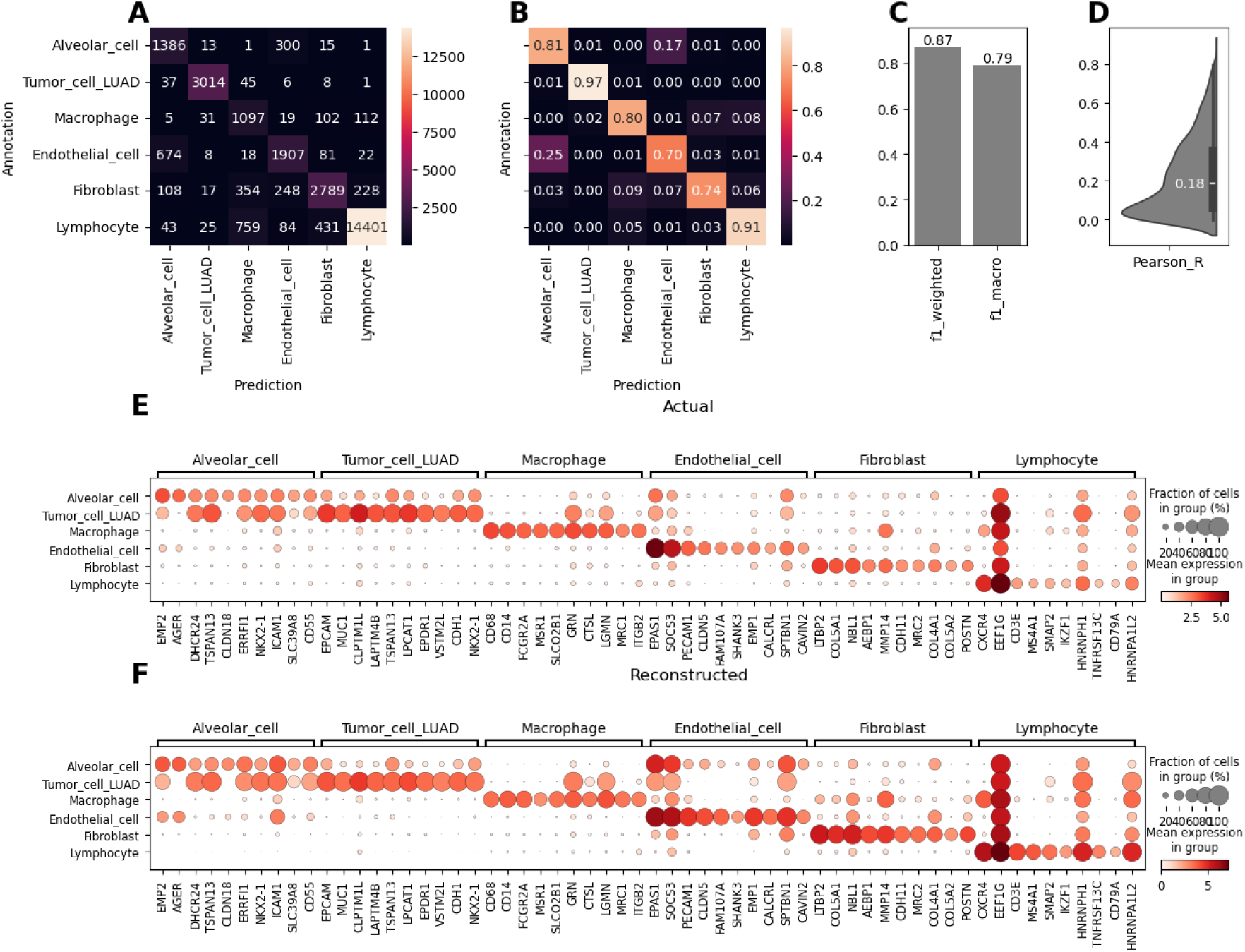
Performance evaluation of the SHEST model on the test dataset derived from the 10X Xenium Prime Human Lung Cancer cohort. **A** Confusion matrix of predicted versus actual cell types, showing the number of correctly and incorrectly classified cells across six cell types. **B** Normalised confusion matrix displaying classification accuracy per cell type. **C** Weighted and macro F_1_-scores of the classifier. **D** Violin plot of the Pearson R distribution of the actual and reconstructed expression of each gene across all cells. **E** Dot plot of marker gene expression in actual data. Dot size represents the fraction of cells expressing each gene, and colour intensity indicates mean expression levels within each cell type. **F** Corresponding dot plot from reconstructed gene expression.

To evaluate the fidelity of the multi-task model in predicting transcriptomic profiles, we first examined the correlation distribution between the actual and reconstructed expression of all genes, which showed that most values exhibited a positive correlation (Figure 4D). This confirms that morphological features encoded by SHEST convey sufficient information to infer functional molecular states.

Subsequently, the top 10 marker genes for each cell type were identified from the ranked gene groups using the actual gene expression data (Figure 4E). The reconstructed expression profile (Figure 4F) was then compared with the actual profile for these selected markers. The results demonstrated high reconstruction fidelity, particularly in reproducing the expression patterns, where the distinct marker gene signatures of each cell type were preserved. The consistency of these patterns indicates that the model effectively captures the morphological determinants of cell identity embedded in histological features, enabling cross-modality translation from image to transcriptome space.

### 3.2 The SHEST model demonstrates external validity in both cell type identification and spatial organisation

To evaluate the generalisability of SHEST, the trained model was applied to two independent Xenium V1 Human Lung Cancer datasets, which differ in gene panel composition and tissue characteristics, in order to assess the transferability of its learnt morphological-molecular representations across distinct experimental and biological environments (Figure 5 and Supplementary Figure S1). Cells located mainly at the tissue periphery, where the H&E image boundaries did not align with the nuclear boundaries in the ST data, were excluded from inference. These correspond to cells with a mean grey value exceeding 230 in their grayscale representation.

**Figure 5:**
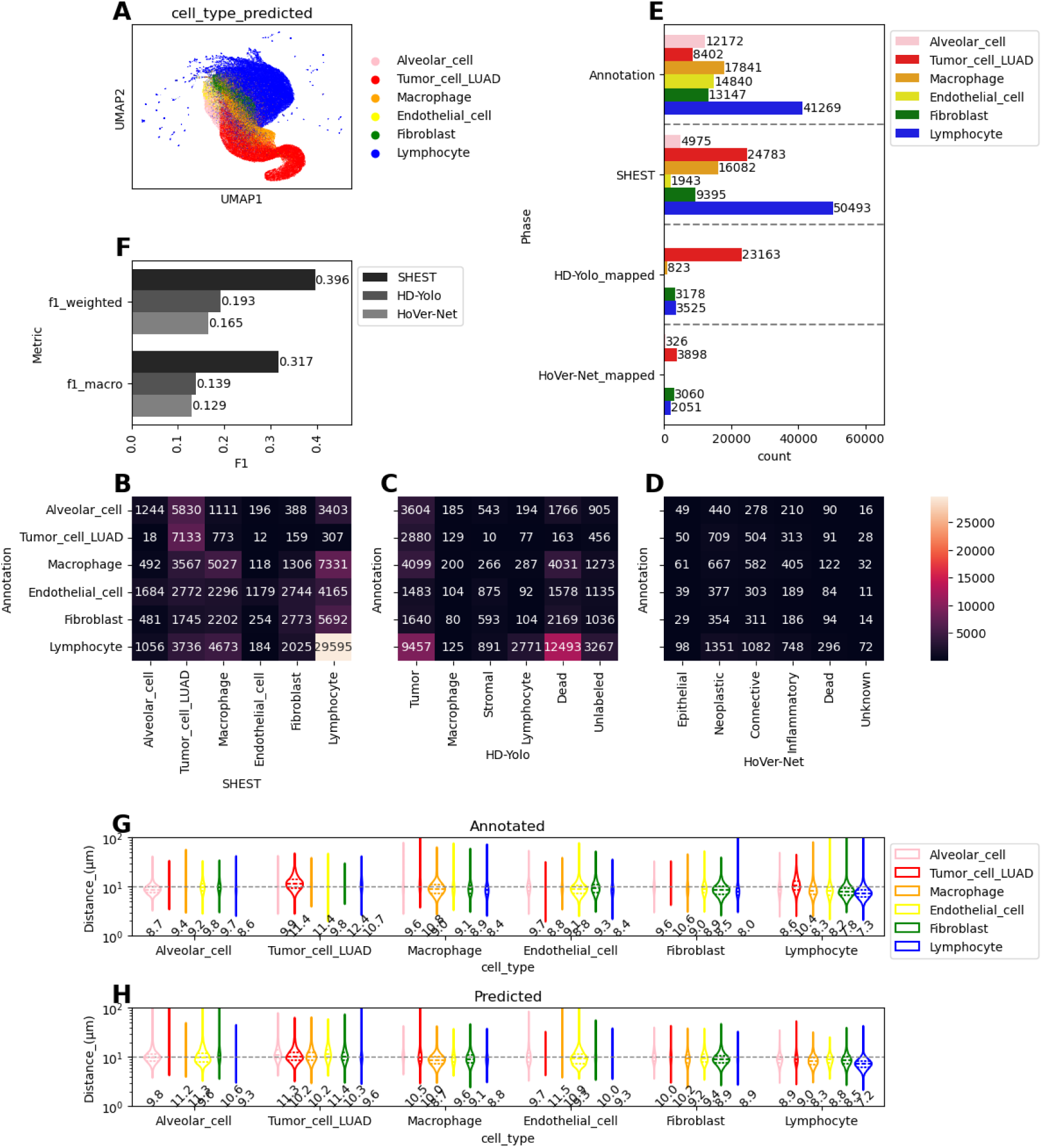
Cell type prediction by the SHEST model on the validation dataset derived from a 10X Xenium V1 Human Lung Cancer Addon panel case. **A** UMAP embedding from the image encoder. **B**-**D** Confusion matrices of predicted cell types, comparing the reference annotation to the predic-tions from SHEST (**B**), HD-Yolo (**C**), and HoVer-Net (**D**) models. **E** Cell type count comparison between annotations and the models. From HD-Yolo and HoVer-Net, cell types were mapped to the category most similar to the reference. **F** W28eighted and macro F_1_ scores of the classification results. **G**-**H** Comparison of spatial cell-cell distances between annotated and predicted cell types. Violin plots showing the distribution of distances from each cell type to its nearest neighbour. The numbers below represent the median. **G** Distances derived from annotated cell types. **H** Distances derived from model-predicted cell types.

The UMAP embedding generated by the image encoder (Figure 5A) demonstrates the model’s ability to cluster cells based on image features while preserving distinct cellular identities. This indicates that the morphological features learnt from the Xenium Prime samples are sufficiently generic to capture cross-sample variability while preserving lineage-specific characteristics. When comparing the annotated cell types with predictions from SHEST, HD-Yolo, and HoVer-Net (Figures 5B–D), SHEST shows superior concordance with the reference annotations (Figure 5B). This observation is further supported by the mapped cell type counts (Figure 5E), indicating that SHEST provides more comprehensive and detailed predictions. In the cross-comparison of *F*_1_ scores (Figure 5F), SHEST achieved the highest performance in both weighted and macro *F*_1_ metrics.

In addition to cell-level accuracy, we examined whether SHEST can preserve spatial organisation at the tissue scale. To assess the fidelity of the spatial organisation inferred by the model, cell–cell distances were compared between the annotated and predicted cell types (Figures 5G and H). The areas of distribution are comparable, and notably, the median distances derived from the annotated data (Figure 5G) and predicted data (Figure 5H) show consistent agreement across all cell type combinations. Particularly, the close clustering of lymphocytes—a hallmark of tertiary lymphoid structure (TLS) formation within the TME [35]—was faithfully recapitulated. This alignment between predicted and actual cell–cell proximities implies that the model’s predictions are not indiscriminate classifications but are spatially coherent in a manner consistent with biological tissue organisation.

We examined the morphological characteristics of cases misclassified as lymphocytes by SHEST (Supplementary Figure S2). The analysis revealed that Macrophages, which are originally large and pale, were misclassified when they appeared smaller and darker; Endothelial_cells, typically located near red blood cells (RBCs), were misclassified when they were positioned away from RBCs and exhibited smaller nuclei; and Fibroblasts, which are normally elongated, were misclassified when they appeared round and small. These observations substantiate that the encoder’s latent space encodes biologically stable morphological cues rather than cohort-specific visual artefacts.

Another noteworthy observation arises when examining the predicted cell maps with tissue histology. In the normal alveolar regions, the model correctly classifies the cells as Alveolar_cell (Figure 6B), whereas within lepidic-pattern areas, indicative of early adenocarcinomatous transformation, it identifies alveolar cells as Tumor_cell_LUAD based on morphological features (Figure 6E). This spatial concordance demonstrates that the model captures histopathologically meaningful transitions, effectively linking structural changes in the tissue to shifts in cellular identity.

**Figure 6:**
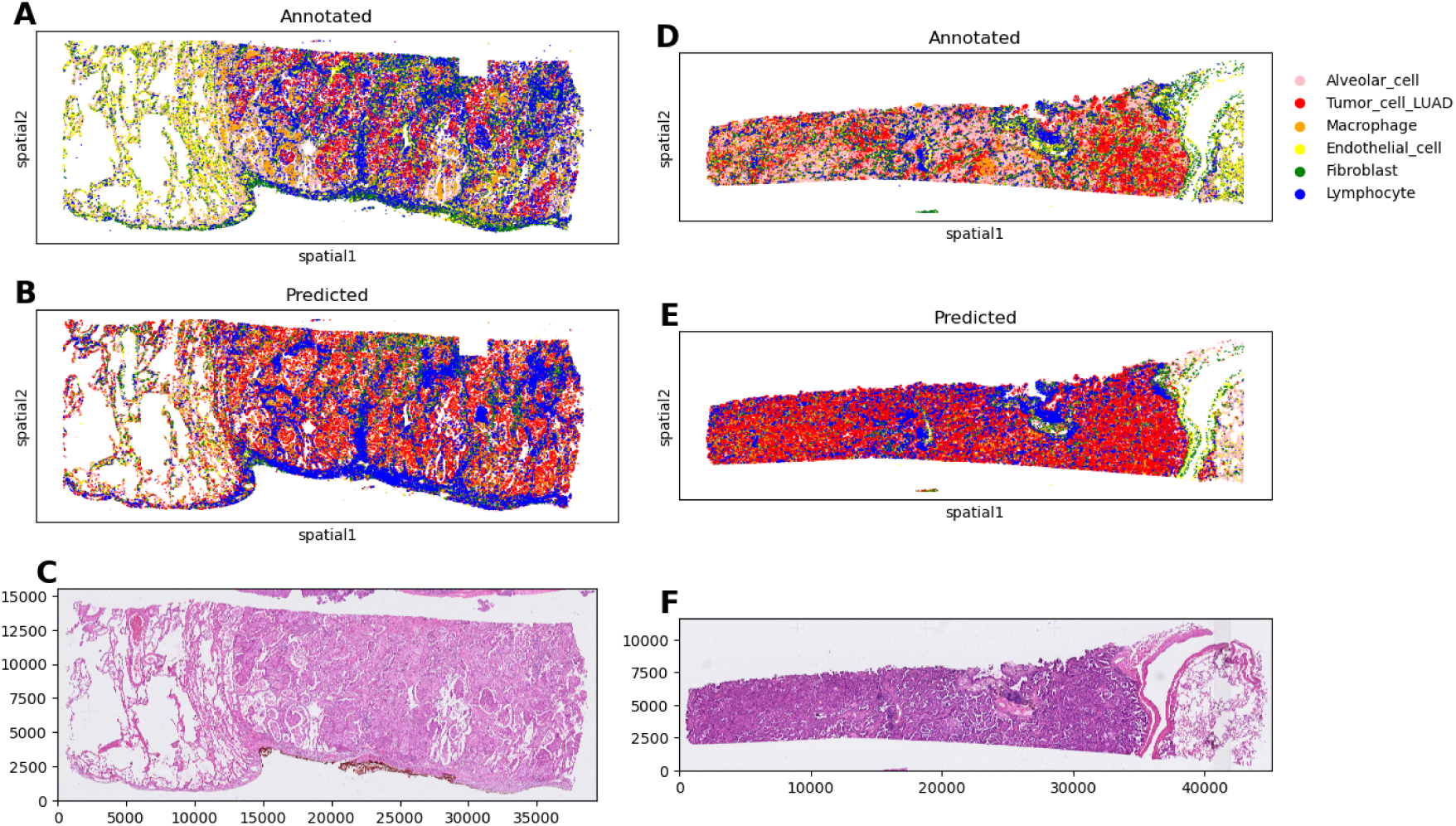
Comparison of annotated and predicted cell types in Xenium V1 Human Lung Cancer datasets with corresponding histology. **A**-**C** Results from the sample with Addon panel: (**A**) annotated cell types by the LuCA single-cell reference, (**B**) predicted cell types by the SHEST model, and (**C**) H&E staining image providing morphological context. **D**-**F** Results from the sample with Multitissue panel: (**D**) annotated cell types by the LuCA single-cell reference, (**E**) predicted cell types by the SHEST model, and (**F**) H&E staining image providing morphological context.

Collectively, these results confirm that SHEST exhibits strong external validity in both cell-type recognition and spatial context preservation. The model retains interpretability and spatial coherence even across different material sources and gene panels, proving that the learnt morphology–molecular correspondence reflects intrinsic cellular biology.

### 3.3 The SHEST model reconstructs not only cell-level gene expression but also global expression patterns from H&E images

The capability of the SHEST model to accurately reconstruct cell-level transcriptomic profiles from histological images is validated through an external validity assessment by comparing the actual and predicted gene expression data (Figure 7 and Supplementary Figure S3).

**Figure 7:**
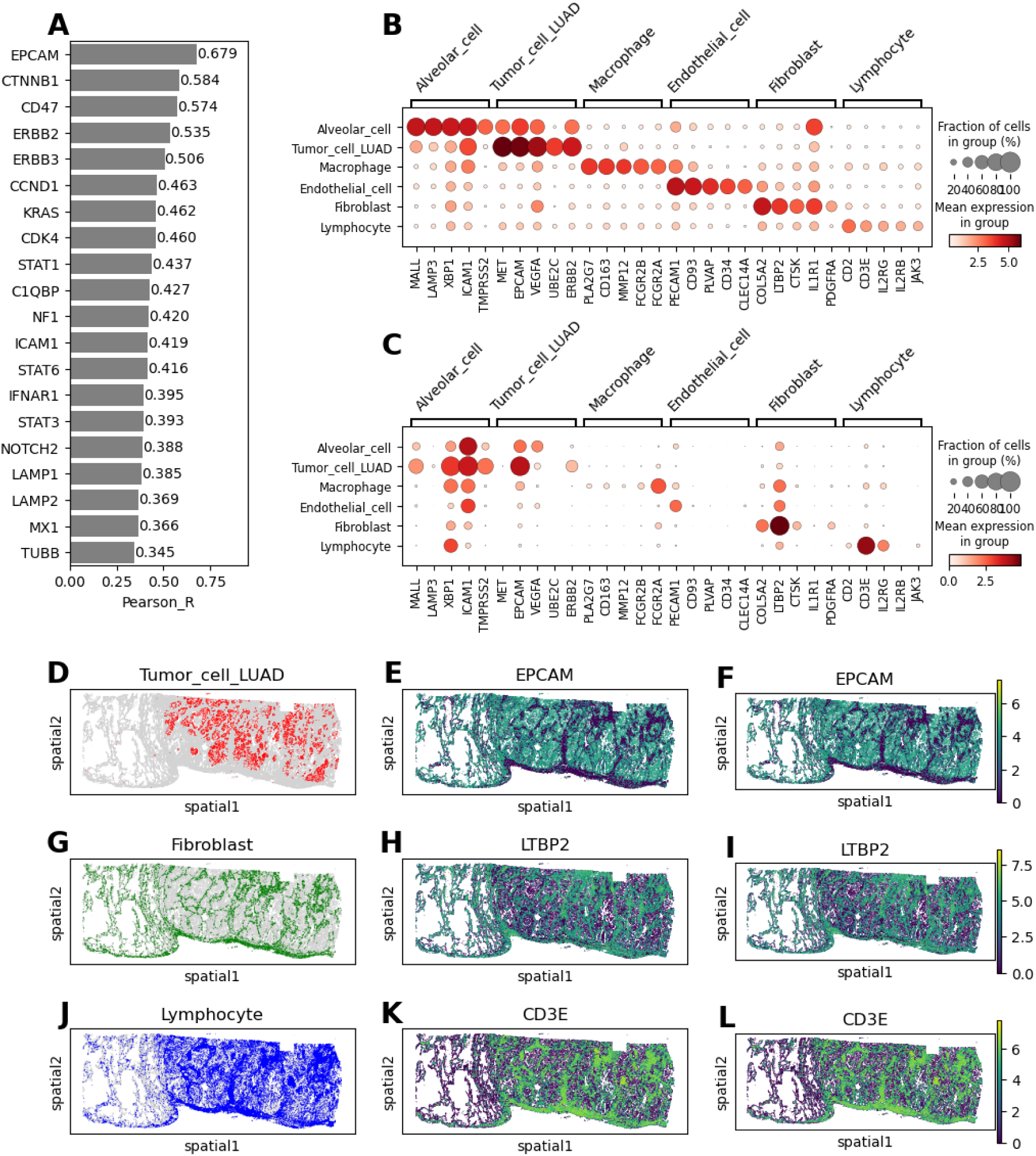
Spatial expression reconstruction by the SHEST model on the validation dataset derived from the 10X Xenium V1 Human Lung Cancer Addon panel case. **A** Top 20 Pearson correlation coefficients for genes comparing actual and reconstructed expression across all cells. **B** Distribu-tion of actual expression for five key marker genes per cell type. **C** Distribution of reconstructed expression for the same marker genes shown in **B**, demonstrating the model’s fidelity in capturing cell-type-specific signatures. **D** Spatial distributi^3^o^0^n map of Tumor_cell_LUAD cells (red) across the tissue section. **E** and **F** Spatial expression map of the tumour marker gene *EPCAM*: actual (**E**) and reconstructed (**F**). **G** Spatial distribution map of Fibroblasts (green). **H** and **I** Spatial expression map of the fibroblast marker gene *LTBP2*: actual (**H**) and reconstructed (**I**). **J** Spatial distribu-tion map of Lymphocytes (blue). **K** and **L** Spatial expression map of the lymphocyte marker gene *CD3E*: actual (**K**) and reconstructed (**L**).

We ranked genes by Pearson correlation between observed and reconstructed single-cell expression and found canonical marker genes among the top-correlating set (Figure 7A). In particular, the tumour epithelial marker *EPCAM*, the fibroblast-associated extracellular matrix gene *LTBP2*, and the lymphocyte marker *CD3E* showed high reconstruction fidelity (Figure 7B and C). In further detail, a comparison of the actual expression (Figure 7B) and reconstructed expression (Figure 7C) for the top five marker genes per cell type confirms that the prominent expression of these marker genes is maintained in both profiles. *EPCAM* reflects the progression and metastasis of cancer in epithelial-derived malignancies [36]. *LTBP2*, secreted by activated fibroblasts and cancer-associated fibroblasts, embodies extracellular matrix remodelling and stromal activation processes essential to tumour progression [37]. *CD3E* defines T-cell identity and immune activation within TLS [38]. These results indicate that the image-derived embeddings are sufficient to recover both lineage identity and quantitative marker expression at the single-cell level, capturing the functional convergence between cell morphology and molecular phenotype.

Another noteworthy point is that SHEST was trained using the more comprehensive Xenium Prime dataset encompassing approximately 5K genes, thereby enabling expression profiling of genes such as *LTBP2* that are not included in the Xenium V1 validation panel. This expanded molecular scope demonstrates the model’s ability to generalise beyond the directly measured transcriptomic range, allowing the inference of biologically relevant genes absent from the validation dataset and supporting a more integrative and mechanistically interpretable reconstruction of the TME.

Beyond single-cell reconstruction, SHEST reproduced global gene-expression landscapes that closely matched the actual data. Spatial maps of reconstructed marker expression aligned well with the actual distributions across tissue sections (Figures 7D–L). In detail, both the actual and reconstructed profiles consistently showed that *EPCAM* expression was concentrated within tumour nests, *LTBP2* delineated fibroblast-rich stromal regions, and *CD3E* was specifically expressed within peritumoral TLS. The high accuracy of this spatial expression reconstruction validates the model’s utility as a tool for generating synthetic ST data directly from histology. Notably, alveolar cells exhibiting a lepidic pattern, although not annotated as tumour cells, also demonstrated that both the actual measurements (Figure S3E) and the reconstructed spatial profiles (Figure S3F) consistently exhibit high *EPCAM* expression. These observations further suggest that the reconstruction framework not only reliably predicts gene expression at single-cell resolution but also preserves subtle microanatomical patterns, enabling the detection of tumour cells that might otherwise be misclassified or overlooked.

Preservation of the inherent spatial structure of gene expression was further assessed using Moran’s *I* spatial autocorrelation (Figure 8 and Supplementary Figure S4). The spatial correlation of key marker genes essential for defining the TME is notable (Figure 8B and Supplementary Figure S4B). This preservation suggests that gene expression reconstruction from histomorphological images by SHEST achieves a high spatial correlation with actual data, indicating that the model captures not only the molecular identity of individual cells but also the multicellular niche architecture based on spatially reconstructed transcriptomics within the tissue.

**Figure 8:**
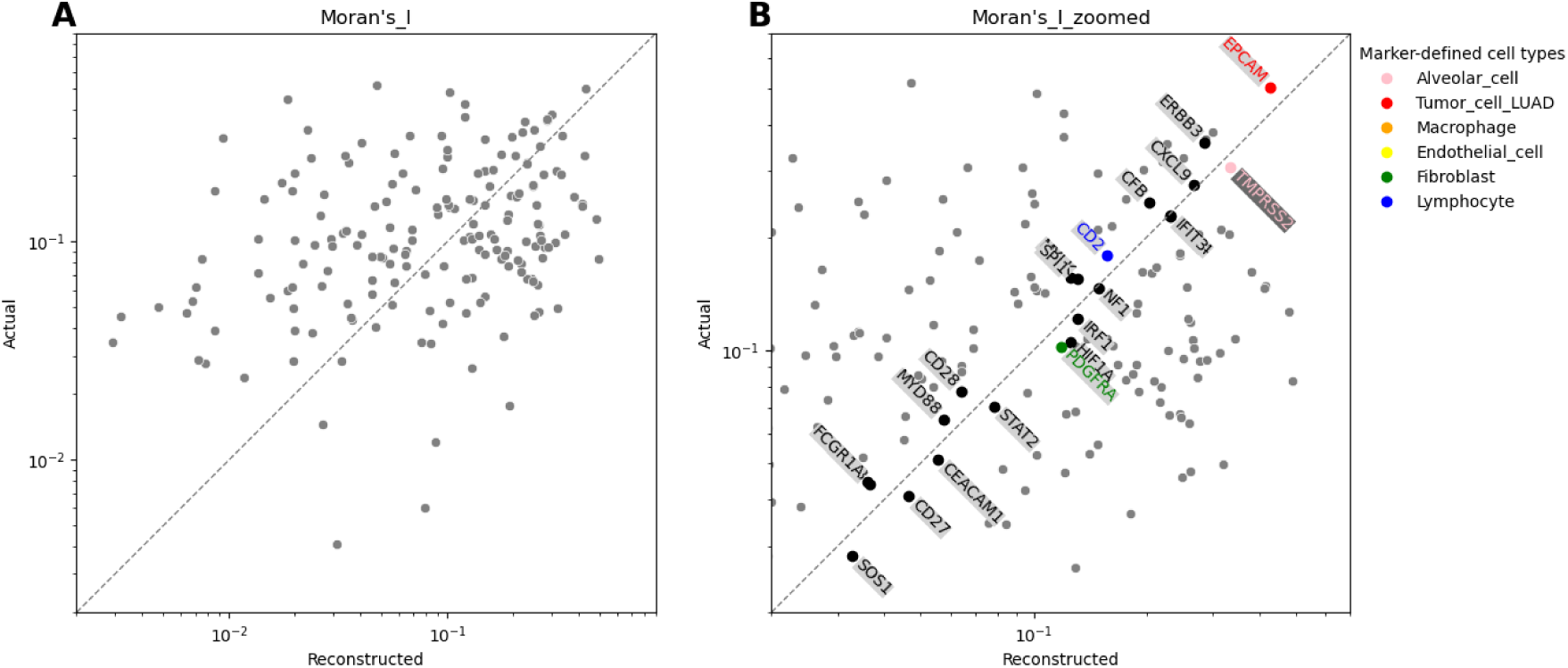
Scatter plot of actual versus reconstructed Moran’s I spatial correlation for individual genes, based on expression values reconstructed by the SHEST model from the Xenium V1 Human Lung Cancer Addon panel dataset. **A** A scatter plot showing the actual versus reconstructed Moran’s I values across the full range. **B** A zoomed-in view of the dense region, with only genes within a 20% relative error margin of the line of identity labelled. Marker genes are colour-coded according to their associated cell type.

## 4 Discussion

The SHEST framework establishes a direct computational bridge between H&E histology and single-cell ST by simultaneously encoding cellular morphology, classifying cell types, and reconstructing transcriptomic profiles at single-cell resolution. This unified modelling approach demonstrates that spatially coherent molecular surrogates can be derived directly from conventional histological slides, connecting tissue architecture and molecular function.

Methodologically, SHEST introduces three principal breakthroughs. (1) The quadruple-tile input structure integrates four complementary image views—whole-cell, nucleus, haematoxylin extraction, and nucleus-specific hematoxylin—thereby encompassing nuclear detail, cytoplasmic morphology, and contextual extracellular matrix information critical for accurate cellular discrimination. (2) A neighbour-aware, cluster-ing–based refinement algorithm selectively removes ambiguous cells from the latent feature space, increasing data purity and ensuring that model training relies on high-confidence examples. This approach reduces multimodal noise and stabilises the convergence of joint optimisation. (3) A ZINB loss function is employed to match the statistical properties of single-cell expression data, effectively modelling overdispersion and sparsity that are inherent to count-based transcriptomic measurements. Together, these elements enable the framework to maintain biological realism in its morphology-to-expression translation while enhancing performance in both classification and reconstruction tasks.

Building upon the research findings, the SHEST framework offers several insights and implications. (1) The framework demonstrates stable inference performance across diverse staining conditions and gene panels, preserving both accuracy and interpretability. Having been trained on diverse staining conditions and extensive gene panels, the SHEST model complements the inherent functional information of a given tissue morphology while maintaining inference consistency. This can serve as a foundational approach for integrative translational medicine, bridging histopathological interpretation with molecular-level diagnosis. (2) The model’s capacity to extrapolate molecular states from morphology implies that it captures higher-order relationships within the morphology–molecular transformation space. This capacity suggests that SHEST can infer unmeasured or functionally latent features, potentially revealing previously uncharacterised cellular functions or molecular signatures [9]. (3) The reconstructed spatial organisation reflects not only cellular identities but also their proximities and densities, elucidating the topological dynamics of the TME. Such spatial coherence provides a valuable substrate for downstream analyses, including cell–cell interaction inference, immune infiltration mapping, and microenvironmental remodelling studies [39]. (4) Trained on multimodal data where morphological and transcriptional identities are jointly validated, SHEST recognises cell types consistent with both morphological integrity and molecular fidelity. This dual validation enables the discrimination of early neoplastic transitions from benign epithelial states and the characterisation of progressive malignant phenotypes, providing a biologically grounded framework for understanding cellular differentiation trajectories and the morphological manifestations of tumour evolution [40, 41].

Despite its innovative outcomes, several limitations remain in this study. Unlike cell type prediction, gene expression reconstruction could not be benchmarked, as previous studies were trained on Visium patches from breast cancer, whereas our work was trained on Xenium WSI images of LUAD. Our study focuses primarily on LUAD, a tumour type selected for the accessibility of case acquisition and cohort construction. Extending the framework to encompass diverse tumour types and tissue microenvironments represents an important direction for future work. In addition, although the training dataset was derived from the state-of-the-art Xenium 5K platform with single-cell resolution, a full-transcriptome gene panel has not yet been made publicly available [42]. As ST technologies advance toward whole-transcriptome coverage, the proposed model can be further extended to achieve more comprehensive and biologically enriched molecular representations of the tissue microenvironment.

SHEST demonstrates potential beyond mere technological advancement, contributing to personalised patient care and transformative clinical decision-making. In practice, SHEST enables identification of cellular composition within the TME directly from accessible H&E slides, offering the potential to predict immune responses and therapeutic resistance, particularly in the context of immuno-oncology. By computationally substituting high-cost ST with morphological inference, the framework achieves comparable informational value at an estimated cost reduction of approximately 99.8%. Such efficiency could substantially accelerate the adoption of spatially informed precision medicine. Ultimately, SHEST represents a step toward computationally enabled translational medicine, where histological morphology is not only descriptive but also quantitatively connected to molecular function. This framework holds potential to enhance patient outcomes and contribute to the long-term goal of overcoming refractory cancers through integrated and personalised therapeutic approaches.

## 5 Abbreviations

CE: Cross-Entropy
DP: Digital Pathology
FC: Fully Connected
H&E: Haematoxylin and Eosin
LN: Layer Normalisation
LUAD: Lung Adenocarcinoma
LuCA: Lung Cancer Atlas
MLP: Multi-Layer Perceptron
MSE: Mean Squared Error
NLL: Negative Log-Likelihood
RBC: Red Blood Cell
scRNA-seq: Single-cell RNA Sequencing
SHEST: Single-cell-level artificial intelligence from Haematoxylin and Eosin morphology for cell type prediction and Spatial Transcriptomics reconstruction
ST: Spatial Transcriptomics
TME: Tumour Microenvironment
TLS: Tertiary Lymphoid Structures
ViT: Vision Transformer
WSI: Whole-Slide Image
ZINB: Zero-Inflated Negative Binomial

## 6 Data availability

Single-cell reference data were obtained from the CELLxGENE Discover datasets: https://cellxgene.cziscience.com/datasets. Public data on lung cancer samples from Xenium platforms were obtained from the 10X Genomics datasets page: https://www.10xgenomics.com/datasets/.

## 7 Code availability

The code for the SHEST model is maintained in the following GitHub repository: https://github.com/ recognizability/shest.

## 8 Declarations

### 8.2 Ethical approval

This study was approved by the Institutional Review Board (IRB) of Samsung Medical Center (IRB No. 2022-11-132).

### 8.4 Authors contributions

Conceptualisation: H.J.; data curation: H.J., J.O., D.L., J.H.K.; formal analysis: H.J.; funding acquisition: H.J. and Y.C.; investigation: H.J.; methodology: H.J.; project administration: H.J.; resources: H.J. and Y.C.; software: H.J.; supervision: H.J. and Y.C.; validation: H.J., J.O., and Y.C.; visualisation: H.J.; writing — original draft: H.J.; writing — review and editing: all authors.

---

**Supplementary Figure S1:**
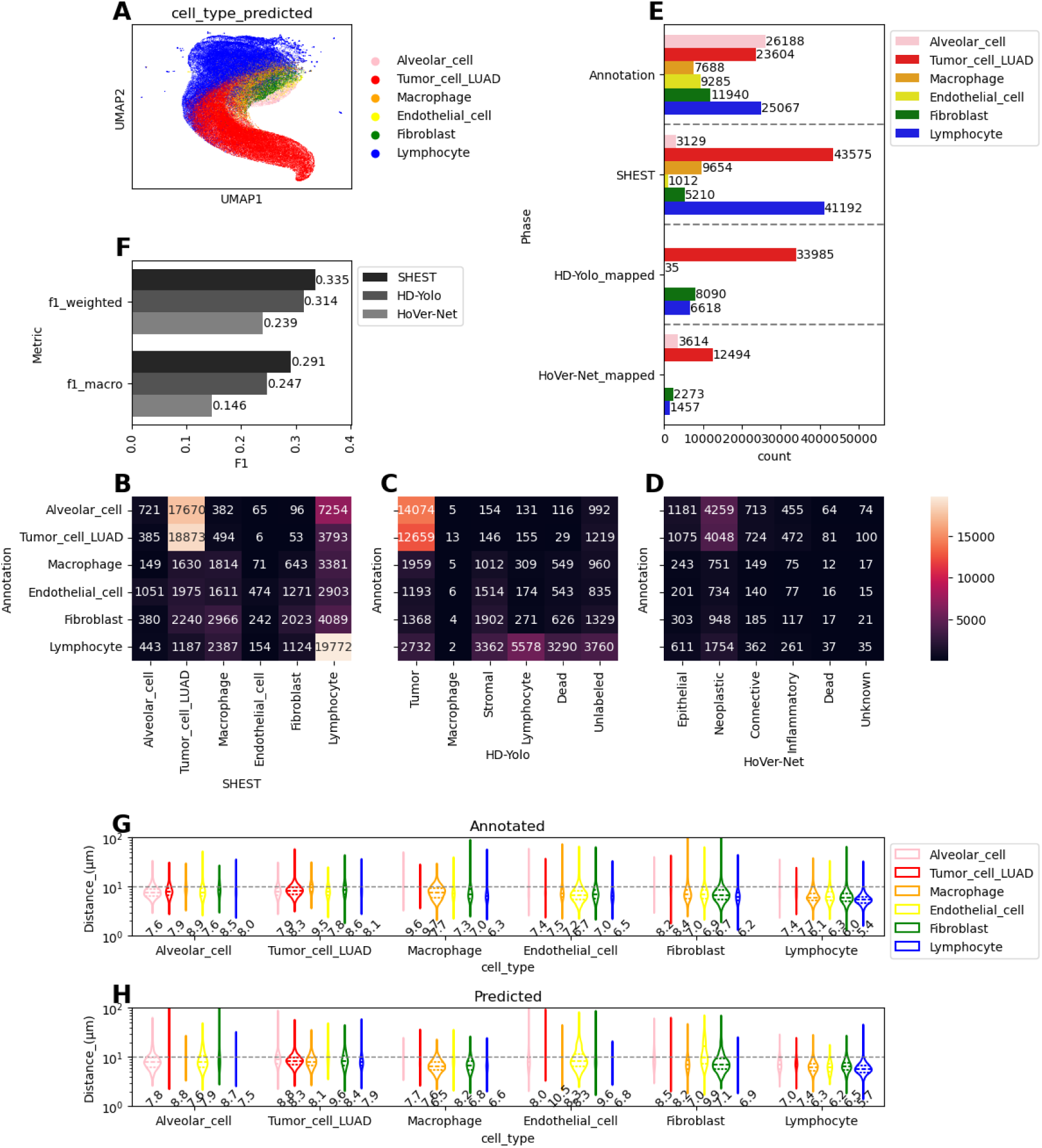
Cell type prediction by the SHEST model on the validation dataset derived from a 10X Xenium V1 Human Lung Cancer Multitissue panel case. **A** UMAP embedding from the image encoder. **B**-**D** Confusion matrices of predicted cell types, comparing the reference annotation to the predictions from SHEST (**B**), HD-Yolo (**C**), and HoVer-Net (**D**) models. **E** Cell type count comparison between annotations and the models. From HD-Yolo and HoVer-Net, cell types were mapped to the category most similar3t2o the reference. **F** Weighted and macro F_1_ scores of the classification results. **G**-**H** Comparison of spatial cell-cell distances between annotated and predicted cell types. Violin plots showing the distribution of distances from each cell type to its nearest neighbour. The numbers below represent the median.**G** Distances derived from annotated cell types. **H** Distances derived from model-predicted cell types.

**Supplementary Figure S2:**
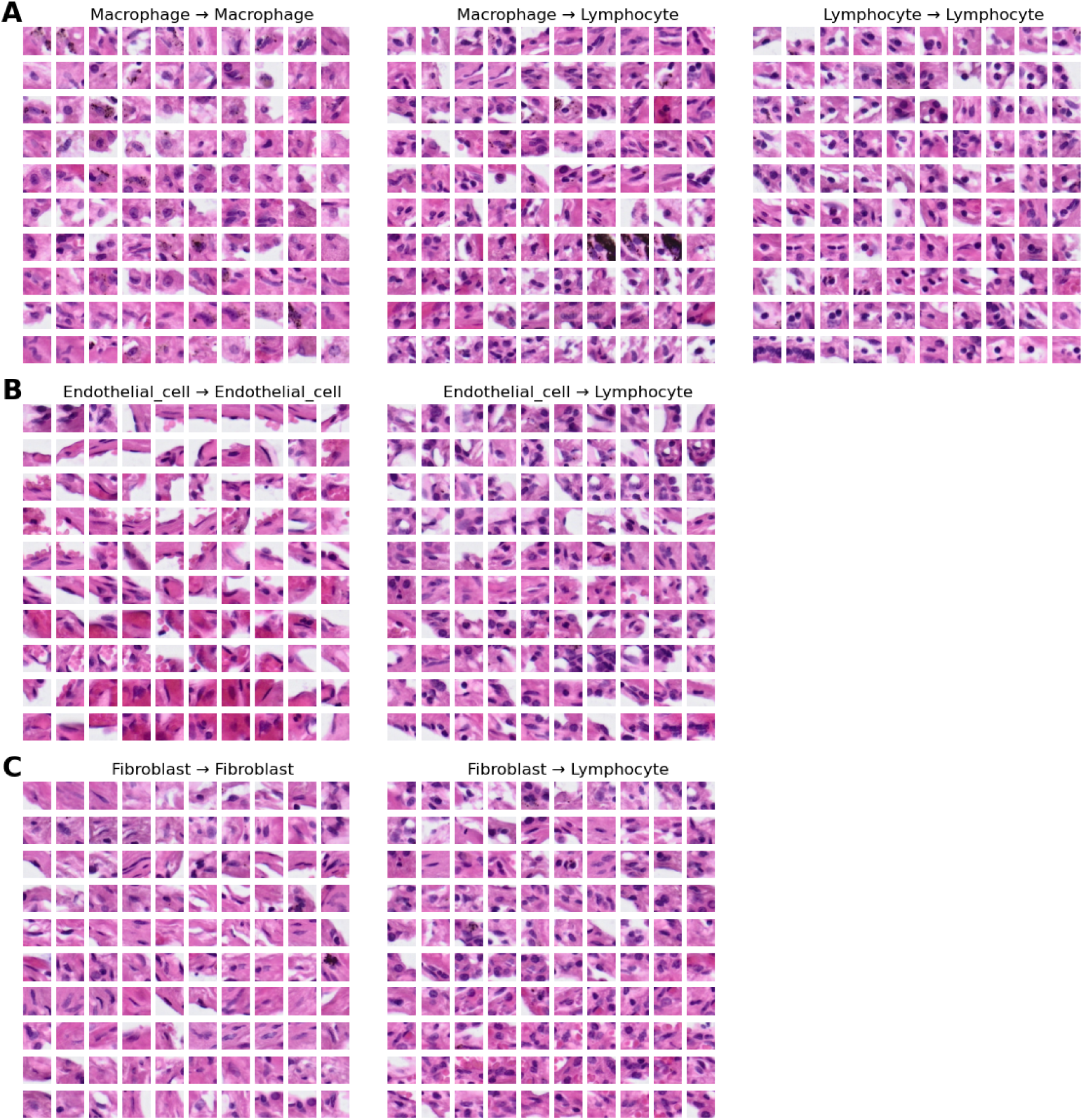
Representative examples of correctly classified and misclassified cases by the SHEST model for the Xenium V1 Human Lung Cancer Addon panel dataset. **A** Macrophage → Macrophage (correct) vs. Macrophage → Lymphocyte (misclassified). The far right is for compar-ison, showing Lymphocyte → Lymphocyte (correct). **B** Endothelial_cell → Endothelial_cell (correct) vs. Endothelial_cell → Lymphocyte (misclassified). **C** Fibroblast → Fibroblast (correct) vs. Fibroblast → Lymphocyte (misclassified).

**Supplementary Figure S3:**
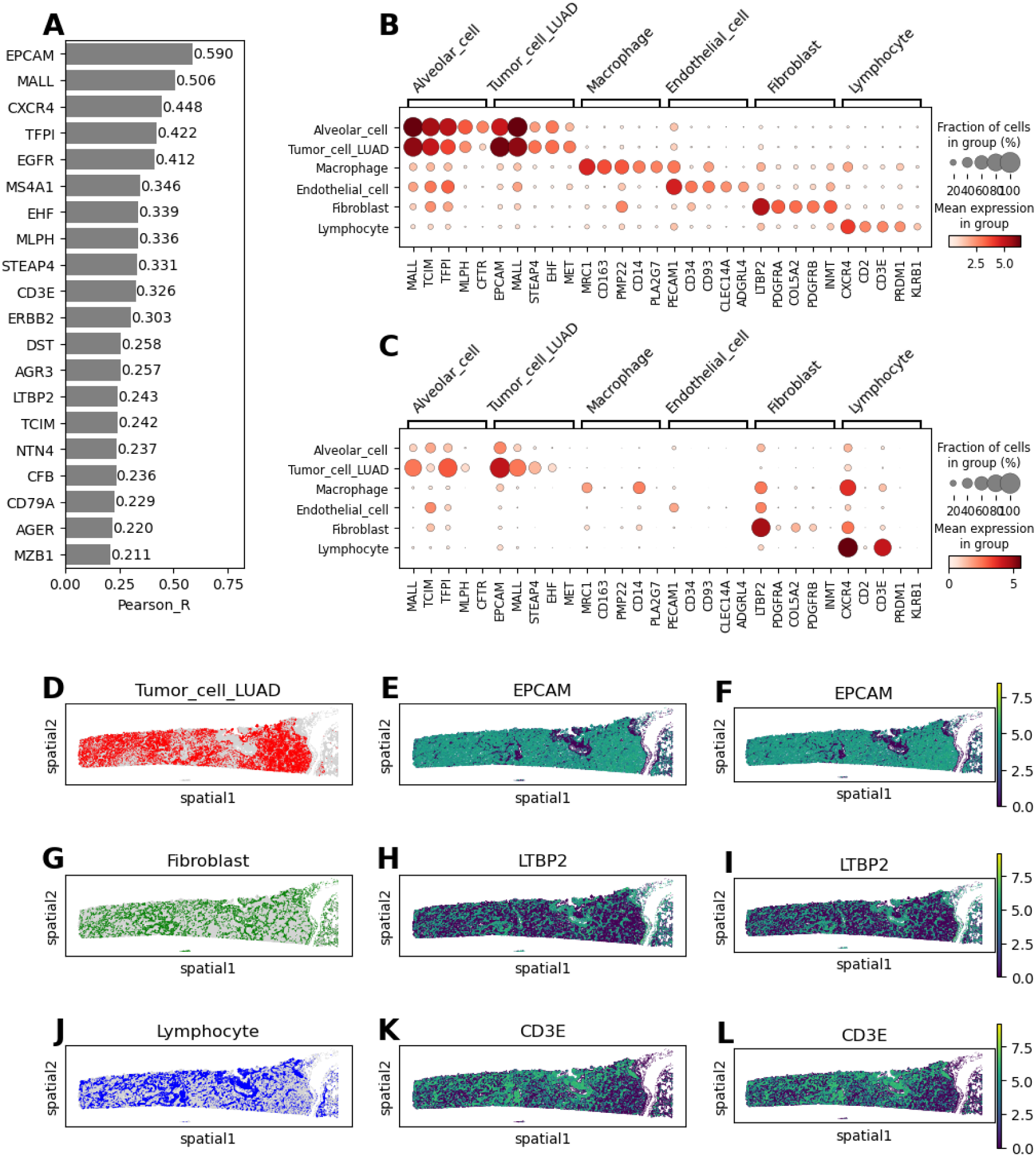
Spatial expression reconstruction by the SHEST model on the validation dataset derived from the 10X Xenium V1 Human Lung Cancer Multitissue panel case. **A** Top 20 Pearson correlation coefficients for genes comparing actual and reconstructed expression across all cells. **B** Distribution of actual expression for five key marker genes per cell type. **C** Distribution of reconstructed expression for the same marker genes shown in **B**, demonstrating the model’s 34 fidelity in capturing cell-type-specific signatures. **D** Spatial distribution map of Tumor_cell_LUAD cells (red) across the tissue section. **E** and **F** Spatial expression map of the tumour marker gene *EPCAM*: actual (**E**) and reconstructed (**F**). **G** Spatial distribution map of Fibroblasts (green). **H** and **I** Spatial expression map of the fibroblast marker gene *LTBP2*: actual (**H**) and reconstructed (**I**). **J** Spatial distribution map of Lymphocytes (blue). **K** and **L** Spatial expression map of the lymphocyte marker gene *CD3E*: actual (**K**) and reconstructed (**L**).

**Supplementary Figure S4:**
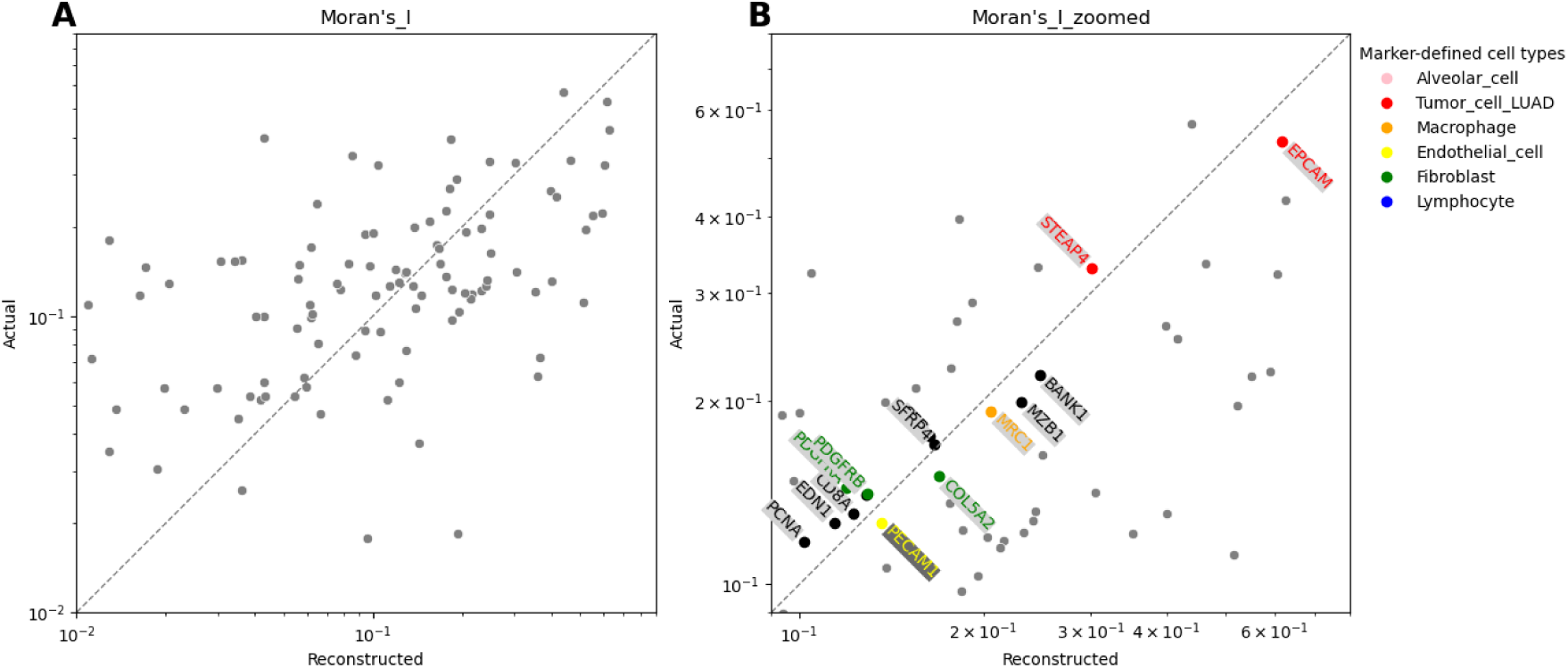
Scatter plot of actual versus reconstructed Moran’s I spatial correlation for individual genes, based on expression values reconstructed by the SHEST model from the Xenium V1 Human Lung Cancer Multitissue panel dataset. **A** A scatter plot showing the actual versus reconstructed Moran’s I values across the full range. **B** A zoomed-in view of the dense region, with only genes within a 20% relative error margin of the line of identity labelled. Marker genes are colour-coded according to their associated cell type.

**Supplementary Table S1:**
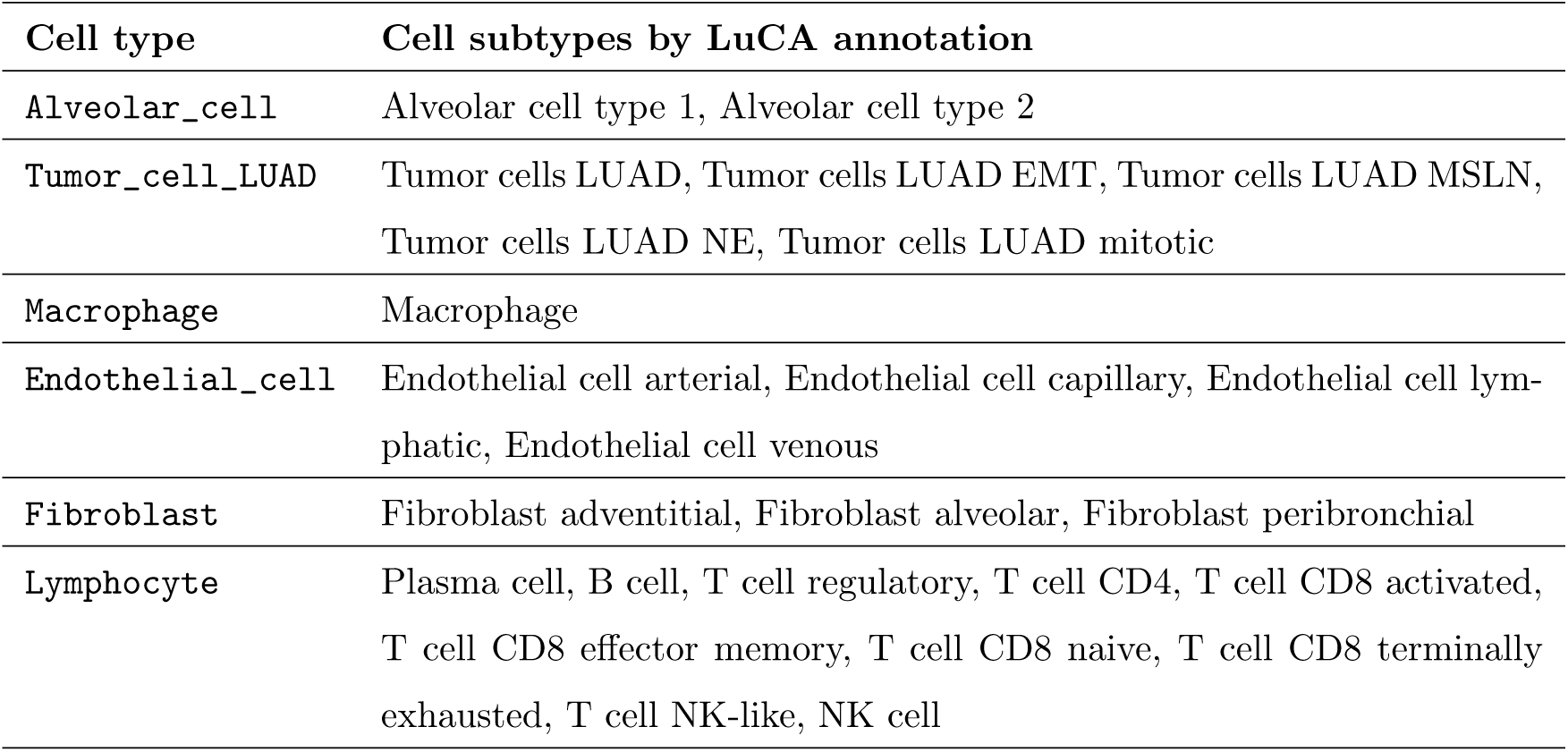
Categorisation of cell subtypes into cell types within the lung cancer tissue.

| **Cell type** | **Cell subtypes by LuCA annotation** |
| --- | --- |
| Alveolar_cell | Alveolar cell type 1, Alveolar cell type 2 |
| Tumor_cell_LUAD | Tumor cells LUAD, Tumor cells LUAD EMT, Tumor cells LUAD MSLN, Tumor cells LUAD NE, Tumor cells LUAD mitotic |
| Macrophage | Macrophage |
| Endothelial_cell | Endothelial cell arterial, Endothelial cell capillary, Endothelial cell lymphatic, Endothelial cell venous |
| Fibroblast | Fibroblast adventitial, Fibroblast alveolar, Fibroblast peribronchial |
| Lymphocyte | Plasma cell, B cell, T cell regulatory, T cell CD4, T cell CD8 activated, T cell CD8 effector memory, T cell CD8 naive, T cell CD8 terminally exhausted, T cell NK-like, NK cell |

